# Mammalian chemosensory bile acid detection supports species and gut microbiome evaluation

**DOI:** 10.1101/2024.10.04.616633

**Authors:** Varun Haran, Mari Morimoto, Leena S. F. Rouyer, Jeffrey G. McDonald, Jinxin Wang, Julian P. Meeks

## Abstract

The rodent accessory olfactory system (AOS) detects environmental chemosignals and guides social and survival-oriented behaviors. Fecal bile acids activate neurons in the AOS, potentially serving as mammalian pheromones and kairomones, but few molecules in this large class have been evaluated thus far. We used live volumetric Ca^2+^ imaging to screen naturally occurring bile acids for their capacity to activate peripheral vomeronasal sensory neurons (VSNs). We found that taurine-conjugated bile acids (tauro-BAs), including taurine-conjugates of cholic acid, deoxycholic acid, lithocholic acid, and chenodeoxycholic acid (TCA, TDCA, TLCA, TCDCA, respectively) activate large populations of VSNs at sub-micromolar concentrations. Tauro-BA-sensitive VSNs rarely responded to unconjugated (CA, CDCA, DCA, LCA), glycine-conjugated (GCA, GDCA, GLCA, GCDCA), or keto-conjugated (7-keto DCA, 12-keto DCA, 7-keto LCA) bile acids. Tauro-BA-sensitive VSNs were also insensitive to well-studied sulfated steroids, suggesting tauro-BAs activate vomeronasal receptors that have not yet been de-orphaned. Among the tauro-BAs, TDCA displayed particularly strong potency, activating many VSNs at sub-micromolar concentrations. Tauro-BAs were not detectable in mouse fecal extracts by mass spectrometry, but were found in reptile fecal extracts and germ-free mouse fecal extracts. VSN responses to germ-free and conventional mouse fecal extracts, conjugated bile acids, and tauro-BAs revealed that non-overlapping populations of VSNs respond to germ-free and conventional mouse feces. A subset of VSNs that were activated by germ-free mouse fecal extracts responded to tauro-BAs, whereas VSNs responsive to conventionally fecal extracts responded to unconjugated bile acids. *In vivo* exposure to TDCA alone, and to mouse fecal extracts spiked with TDCA, elicited mild aversion and stress-associated behaviors in a non-social context (avoidance, digging, grooming, etc.). These studies establish tauro-BAs as a novel class of aversive vomeronasal ligands that vary in feces across species and gut microbiomes.

**Short abstract:** The rodent accessory olfactory system (AOS) detects environmental chemosignals and guides social and survival-oriented behaviors. Fecal bile acids activate neurons in the AOS, but the full repertoire of bile acids that activate VSNs is not known. A live Ca^2+^ imaging screen revealed that VSNs were highly activated by taurine-conjugated bile acids (tauro-BAs) at sub-micromolar concentrations. Tauro-BAs were undetectable in conventionally raised mouse fecal extracts, but were found in reptile fecal extracts and germ-free mouse fecal extracts. VSN activity to tauro-BAs overlapped that of germ-free mouse fecal extracts, suggesting a link between bile acid chemosensation and the gut microbiome. When mice were exposed to taurodeoxycholic acid *in vivo*, they displayed mild aversion and stress-associated behaviors.

## Introduction

Terrestrial mammals, including mice, use multiple olfactory pathways to process chemosensory information, including the main olfactory system (MOS) and the accessory olfactory system (AOS) (Dulac, et al., 2003; Keller, et al., 2009; Suárez, et al., 2012). The MOS consists of the main olfactory epithelium (MOE), which senses volatile environmental odors before relaying this information to the main olfactory bulb (MOB) and downstream brain regions. The AOS, on the other hand, processes social chemosensory information such as pheromones, which are initially detected by the vomeronasal organ (VNO). This information is then sent to the accessory olfactory bulb (AOB) and downstream brain regions. The AOS is important for many different behaviors, such as mating, predator avoidance, and territorial aggression (Mohrhardt, et al., 2018).

A major obstacle to understanding odor processing in the AOS is that only a few ligands (chemosignals) and their corresponding receptors are known for this system. Fecal bile acids (BAs) have previously been found in our lab to function as natural ligands for the mouse AOS (Doyle, et al., 2016; Wong, et al., 2020). Bile acids are essential polar steroids that represent the primary pathway of cholesterol metabolism and play a crucial role in the absorption of lipids and vitamins (Choudhuri, et al., 2022). Analysis of VSN tuning to common BAs using volumetric Ca^2+^ imaging of GCaMP6s revealed multiple populations of BA-responsive VSNs with submicromolar sensitivities (Wong, et al., 2020). However, the extent to which the AOS differentiates and encodes BA chemosensory information at the peripheral level is still uncertain. Additionally, it is also not clear how BA signals are integrated and transformed into codes that guide mouse behavior. Using the AOS as a model system, we hope to understand how sensory neural circuits have evolved to efficiently extract information from the environment that can be used to control “mission-critical” decisions and behaviors.

Primary bile acids, such as cholic acid (CA) and chenodeoxycholic acid (CDCA), are produced in the liver and then converted into the secondary bile acids deoxycholic acid (DCA) and lithocholic acid (LCA), respectively, in the gut through gut microbial metabolism. Each of these bile acids can be conjugated to taurine and glycine, aiding in the digestive process, and facilitating their recycling through the hepato-biliary system (Ramirez-Perez, et al., 2017). Recent reports suggest that there are thousands of potential modifications of bile acids, and that microbes play an essential role in bile acid diversification (Mohanty et al., 2024). How an animal’s age, sex, diet, and intestinal flora regulate the secretion of chemosensory bile acids is not fully known. It is also not known whether gut microbial metabolism modulates bile acid chemosensation in ways that guide the recipient animal’s behavior.

Here, we present data showing taurine-conjugated bile acids (tauro-BAs) are potent VSN ligands. A minority of VSNs activated by tauro-BAs were also sensitive to sulfated steroids, while free amino acids were minimally active, even at high concentrations. Tauro-BAs were undetectable in mouse fecal extracts, but were detected in reptile fecal extracts and germ-free mouse fecal extracts, linking tauro-BA excretion to species, and gut microbiome. Germ-free mouse fecal extracts showed an almost non-overlapping VSN activation pattern compared to conventional fecal extract, and many tauro-BA-responsive VSNs also responded to germ-free mouse fecal extracts. Among the tauro-BAs, taurodeoxycholic acid (TDCA) displayed especially strong potency, activating many VSNs at sub-micromolar concentrations that were unresponsive to other BAs. To test the behavioral impact of tauro-BAs, we exposed mice *in vivo* to TDCA, which induced mild aversion and stress-associated behaviors. These studies identify links between species, gut microbiome, and bile acid chemosensation, and greatly expand our knowledge of AOS function.

## Results

### A calcium imaging screen identifies taurine-conjugated bile acids as potent activators of vomeronasal sensory neurons

Previous studies investigated the sensitivity and selectivity of vomeronasal sensory neurons (VSNs) to common primary (CA, CDCA) and secondary (DCA and LCA) bile acids (Wong, et al., 2020). These studies utilized live volumetric light sheet-based imaging (objective-coupled planar illumination, or OCPI, microscopy), a powerful technique that allows simultaneous monitoring of thousands of VSN per tissue (Holekamp, et al., 2008). The first four characterized bile acids represent a small fraction of the potentially thousands of naturally occurring bile acids ligands that may also serve as AOS chemosignals. We therefore developed and implemented an OCPI-based screen for bile acid VSN ligands. We did so in Omp-Cre x Ai96 animals, which natively express the genetically encoded Ca^2+^ indicator GCaMP6s in VSNs (Li, et al., 2004; Madisen, et al., 2015)(Fig. 1A). All screened stimuli were dissolved in Ringer’s saline solution, and all experiments utilized Ringer’s solution with vehicle controls (< 0.5% methanol) as a negative control. Stimuli were delivered using an established randomized, interleaved block approach that minimizes false positive results (Turaga, et al., 2012)(Figure S1A).

**Figure 1).**
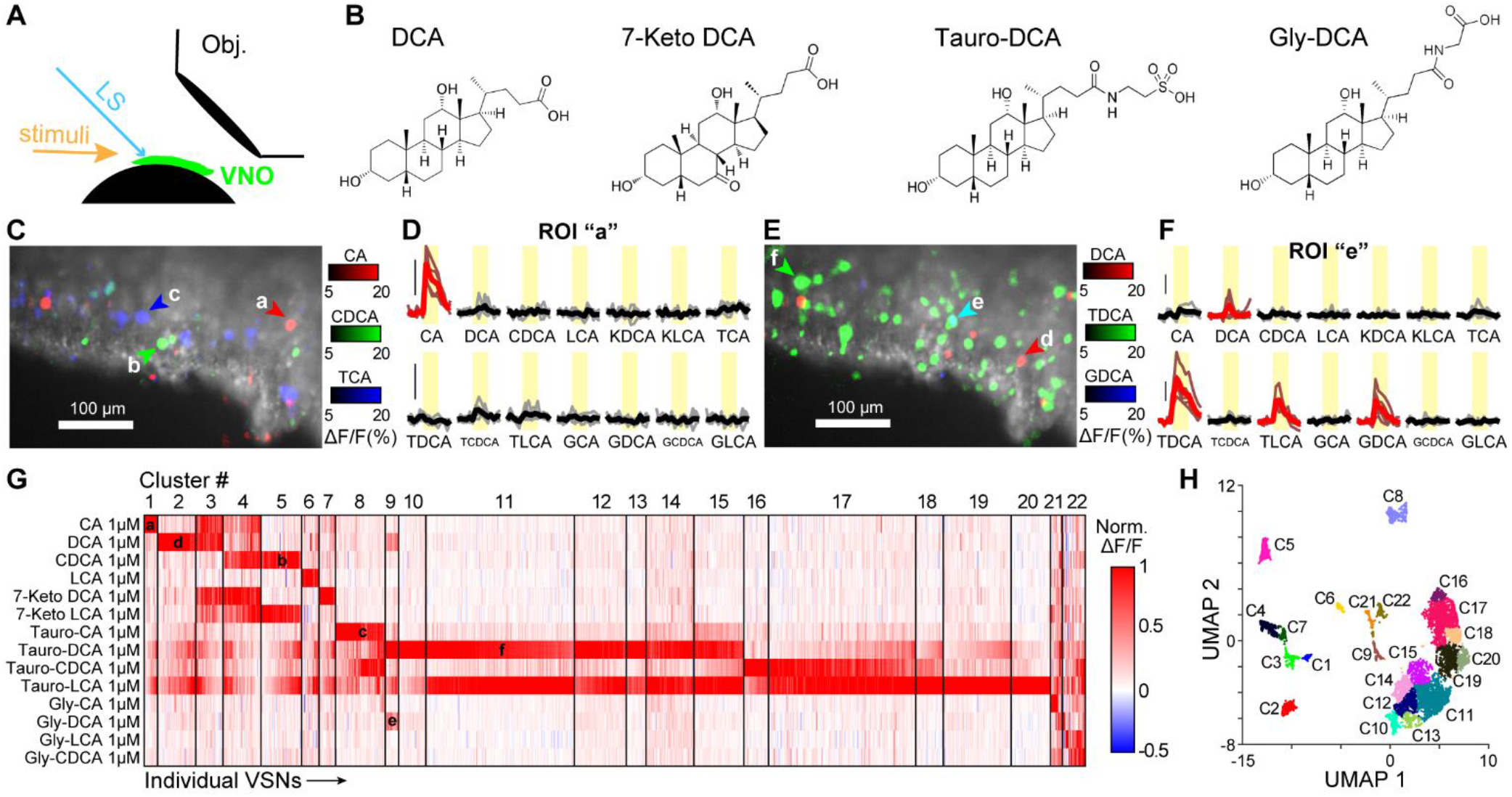
OCPI screen for bile acid ligands (all at 1 µM). (**A**) Diagram of objective-coupled planar illumination (OCPI) microscopy. LS: light sheet. Obj.: objective. (**B**) Chemical structures of deoxycholic acid (DCA) and 7-keto, tauro-, and glyco-conjugates. (**C**) Single image frame from a 51-frame stack showing GCaMP6s-expressing VSNs (gray). Overlaid colors represent ΔF/F responses to CA (red), CDCA (green), and Tauro-CA (blue). Colored arrowheads indicate neurons belonging to the associated clusters in Panel G. (**D**) Peri-stimulus time histograms of VSN stimulus responses. Bold traces reflect mean responses to 5 stimulus repeats. Red traces are statistically larger than control Ringer’s saline (not shown). Yellow rectangles indicate the time of stimulus delivery. Vertical black bars indicate 20% ΔF/F. (**E-F**) Same as Panels C-D but with DCA (red), TDCA (green), and Gly-DCA (GDCA, blue). (**G**) Clustered heat map of ∼8,000 responsive VSNs (11 tissues from 4 males, 3 females). Note the large proportion of VSNs activated by 1 µM taurine-conjugated bile acids. (**H**) UMAP (dimensionality reduction) plot of the tuning properties of the VSNs shown in Panel C. Clusters match Panel G.

Our screen utilized multiple bile acid stimulus panels that were able to compare the sensory profiles of VSNs that were sensitive to established primary and secondary bile acids to those of test ligands, including conjugated bile acids (to taurine, glycine, and the keto modification of bile acids)(Fig. 1B). A full list of the ligands and concentrations screened is presented in Table 1. For each experiment, we manually identified volumetric regions of interest (ROIs) encompassing stimulus-responsive VSNs, and built libraries of VSN responsiveness to screened bile acids ligands, all initially delivered at 1 µM (on the lower end of the concentration range for unconjugated bile acids)(Wong, et al., 2020). Among the screened ligands, we found that tauro-BAs (including TCA, TDCA, TCDCA, and TLCA) activated many VSNs at this concentration. Inspection of the across-ligand tuning properties of tauro-BA-responsive VSNs indicated that most of these VSNs did not respond to other screened ligands (Fig 1C-G). Across multiple tissues from mice of both sexes, we found that the number of VSNs that selectively responded to 1 µM tauro-BAs dwarfed all other screened ligands (Fig. 1G). Notably, tauro-BAs so strongly activated the VNO that some tissues would transiently experience broad/regional fluorescence increases, sometimes causing contamination of ROIs that had no obvious cellular pattern of fluorescence increase (Fig. 1E). This effect can be visualized as horizontal “stripes” in the clustered heatmap in Fig. 1G.

**Table 1.** Mass spectrometry analysis of unconjugated bile acids (CA, DCA, CDCA, LCA), tauro-BAs (TCA, TDCA, TCDCA, TLCA), and glycine-conjugated bile acids (GCA, GDCA, GCDCA, GCLA) in reptile samples. All values normalized to the maximum detected across all animals.

| Genus<br>Common<br>name | Heloderma<br>Beaded<br>lizard | Varanus<br>Perentie<br>monitor | Dendroaspis<br>Green<br>mamba | Shinisaurus<br>crocodile<br>lizard | Phrynosoma<br>horned<br>lizard | Corucia<br>skink | Chelonoidis<br>tortoise | Cyclura<br>iguana |
| --- | --- | --- | --- | --- | --- | --- | --- | --- |
| CA | 0.03 | 0.50 | 0.26 | 1.00 | 0.09 | 0.01 | 0.00 | ---- |
| DCA | 0.26 | 1.00 | 0.23 | 0.06 | 0.60 | 0.03 | 0.01 | 0.13 |
| CDCA | ---- | ---- | ---- | ---- | 0.50 | ---- | ---- | 0.86 |
| LCA | 0.95 | 0.99 | 0.96 | 0.98 | 0.99 | 1.00 | 0.99 | 1.00 |
| TCA | 0.16 | 0.67 | 0.04 | 0.02 | 1.00 | ---- | ---- | ---- |
| TDCA | ---- | 1.00 | ---- | ---- | ---- | ---- | ---- | ---- |
| TCDCA | 1.00 | ---- | ---- | ---- | 0.29 | ---- | ---- | ---- |
| TLCA | ---- | 1.00 | ---- | ---- | 0.62 | ---- | ---- | ---- |
| GCA | ---- | ---- | ---- | ---- | ---- | ---- | ---- | ---- |
| GDCA | 0.02 | 0.09 | ---- | 0.01 | 0.02 | ---- | 0.01 | ---- |
| GCDCA | ---- | 0.19 | ---- | 0.08 | 0.18 | ---- | 0.07 | 0.03 |
| GLCA | 0.89 | 0.90 | 0.93 | 0.76 | 0.90 | 1.00 | 0.88 | 0.96 |

We combining analyzed ROIs from multiple tissues and animals using cluster analysis (see Methods) in order to identify populations of similarly-tuned VSNs (Fig. 1G). Previous studies have linked patterns of VSN ligand sensitivity to cells expressing specific vomeronasal receptors (Haga-Yamanaka, et al., 2014; Lee, et al., 2019; Wong, et al., 2020). We identified 22 distinct clusters, including clusters consistent with prior work showing neurons selective for 1 µM CA (Cluster 1), 1 µM DCA (Cluster 2), and 1 µM LCA (Cluster 6). Also consistent with prior studies, we noted VSN clusters that were tuned to multiple unconjugated bile acids (Clusters 3, 4). These VSNs each showed responsiveness to 7-keto bile acid conjugates 7-keto DCA and 7-keto LCA, expanding the list of known bile acid ligands and extending knowledge about the tuning breadth of BA-sensitive vomeronasal receptors. A small population of VSNs appeared to respond to multiple versions of DCA, including unconjugated DCA, tauro-DCA, and glycine-conjugated DCA (GDCA, Cluster 9). This population, which also responded to Tauro-LCA, indicates the presence of vomeronasal receptors capable of detecting a core bile acid across multiple conjugations. The overwhelmingly largest group of observed VSNs responded to taurine-conjugated BAs (Clusters 8-20). One of these, Cluster 8, contained VSNs that were sensitive to all four screened tauro-BAs, with minimal responsiveness to any other ligand, suggesting the presence of a broad tauro-BA receptor-expressing population. Many tauro-BA-sensitive clusters were nearly identically tuned (e.g., Clusters 11-14), which may be the result of the technical details of cluster analysis (in which parameters must be chosen such that both common and rare subsets are distinguished). Dimensionality reduction methods (UMAP)(McInnes, et al., 2018) showed that the large populations of tauro-BA-sensitive VSNs coalesced into two large macro-clusters (C10-C15 and C16-20) that were clearly separated from all other clusters. Collectively, these data suggest the presence of multiple novel classes of VSNs (and receptors), the largest of which are selective for tauro-BAs.

### Most tauro-BA-sensitive VSNs are insensitive to sulfated steroids

Previous studies identified a subset of VSNs that responded broadly to unconjugated bile acids and sulfated glucocorticoids, likely through the expression of V1R receptors from the “V1re” clade (Isogai, et al., 2011; Wong, et al., 2020). Given the large number of studies into sulfated and other polar steroid responses of VSNs (Fu, et al., 2015; Haga-Yamanaka, et al., 2014; Meeks, et al., 2010; Nodari, et al., 2008; Turaga, et al., 2012; Xu, et al., 2016), we tested whether tauro-BA-responsive VSNs were also sensitive to sulfated steroids (Fig. 2). We performed Ca^2+^ imaging studies in which we exposed the VSNs to the following stimulus conditions: unconjugated bile acids (CA, DCA, CDCA, and LCA; each at 1 μM), tauro-BAs (TCA, TDCA, TLCA, and TCDCA; each at 1 μM), iso-DCA and DHCA (each at 1 μM), and three sulfated steroids (Q1570, A6970, and E0893; each at 10 μM). In this experiment, mouse female feces extract was used as a positive control (not shown).

**Figure 2).**
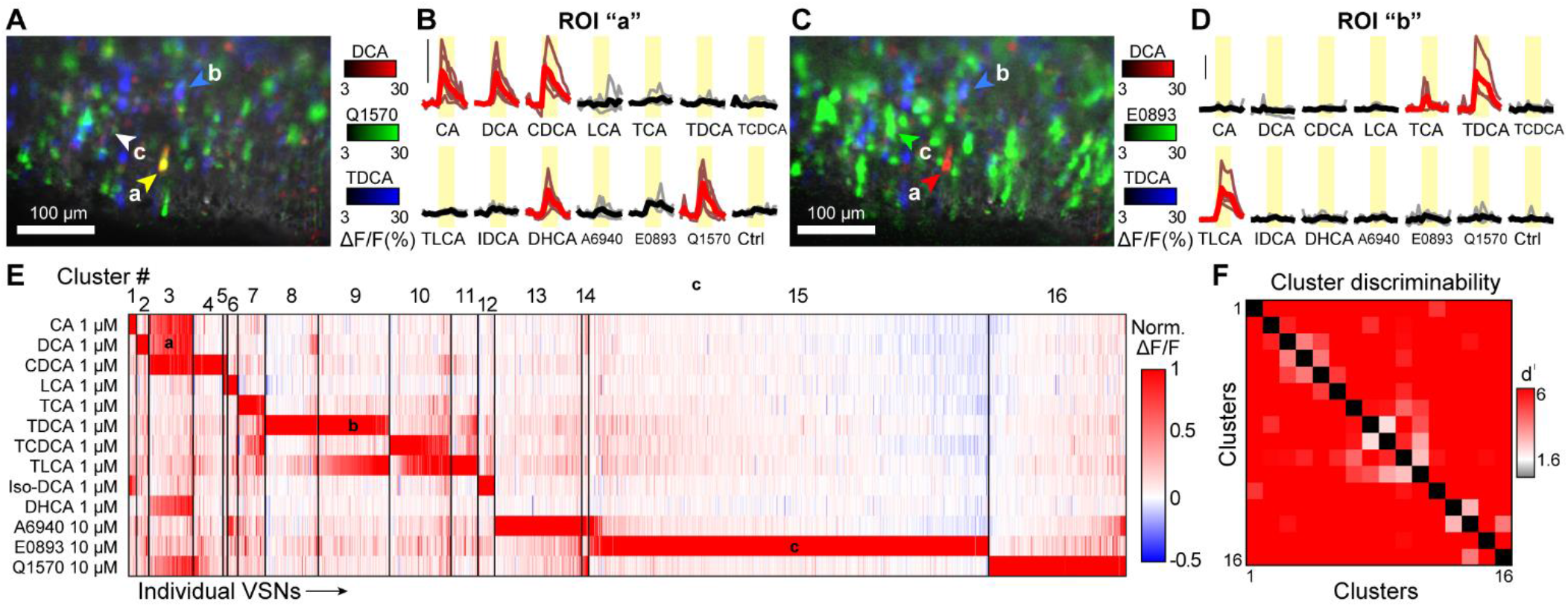
Most tauro-BA-sensitive VSNs do not respond to sulfated steroids. (**A**) Single image frame from a 51-frame stack showing GCaMP6s-expressing VSNs (dim gray). Overlaid colors represent ΔF/F responses to DCA (1 µM, red), sulfated glucocorticoid Q1570 (10 µM, green), and TDCA (1 µM, blue). Arrowheads refer to cluster labels in Panel E. (**B**) Peri-stimulus time histograms of stimulus responses to ROI “a” in Panel A. Bold traces reflect mean responses to 5 stimulus repeats. Red traces are statistically larger than control Ringer’s saline (Ctrl, Wilcoxon Rank Sum Test, p<0.01). Yellow rectangles indicate the time of stimulus delivery. Vertical black bars indicate 40% ΔF/F. (**C**) Overlaid colorized responses from the same image frame as Panel A to DCA (1 µM, red), E0893 (10 µM, green), TDCA (1 µM, blue). Arrowheads refer to cluster labels in Panel E. (**D**) Peri-stimulus time histograms of stimulus responses to ROI “b” in Panel C. All other features same as Panel B. (**E**) Clustered heat map of 5,068 responsive VSNs (9 tissues from 3 males, 3 females). (**F**) Pairwise discriminability index (d^I^) plot for Clusters 1-16 in Panel E. For each comparison, cells in each cluster were projected along the first linear discriminant Eigenvector, and d^I^ calculated from these projections. Red hues represent increasingly statistical separation. White represents a model statistical p-value of 0.05. Gray hues would indicate a lack of statistically significant separation.

Each of the sulfated steroids in this panel – an androgen (A6940), estrogen (E0893), and glucocorticoid (Q1570) – activated specific subsets of VSNs, similar to previous studies (Clusters 13-16, Fig. 2E) (Meeks, et al., 2010; Turaga, et al., 2012; Wong, et al., 2020). Cluster analysis of 5068 VSNs pooled across multiple tissues and male and female animals confirmed the presence of a population of VSNs that were activated by multiple unconjugated bile acids and the sulfated glucocorticoid Q1570 (corticosterone-21-sulfate, 10 µM)(Cluster 3, Figure 2)(Nodari, et al., 2008; Turaga, et al., 2012; Wong, et al., 2020). We did not observe Q1570 sensitivity in tauro-BA-sensitive VSN clusters (Clusters 7-12, Figure 2). Some VSNs that were responsive the sulfated estrogen E0893 (17α-estradiol-3-sulfate, 10 µM), and the sulfate androgen A6940 (epitestosterone 17-sulfate, 10 µM) also responded minimally to tauro-BAs (Clusters 13-15, Fig. 2). There were a relatively small number of these cells, and their across-trial reliability was low, suggesting any overlap in VSN sensitivity between tauro-BAs and sulfated steroids is minimal. This is noteworthy, as the sulfated estrogen E0893, and related sulfated estrogens, are among the most well-studied steroidal VSN ligands (Haga-Yamanaka, et al., 2014; Lee, et al., 2019; Nodari, et al., 2008). Moreover, E0893 and the other steroids were applied at 10 µM, a 10-fold higher concentration than tested bile acids. E0893 was the most active stimulus in this test panel, with cells assigned to Cluster 14 comprising 40% of analyzed cells (1993/5068). Collectively, these data suggest that tauro-BA-sensitive VSNs likely express vomeronasal receptors that have not been “de-orphaned.”

### VSNs are sensitive to taurine-conjugated bile acids at sub-micromolar concentrations

Given the large number of 1 µM tauro-BA-responsive VSNs, we hypothesized that tauro-BAs may activate VSNs at very low concentrations. LCA and derivatives (e.g. TLCA) have previously been associated with acute cellular toxicity (Graf, et al., 2002; Katona, et al., 2009), which may at least partially account for their broad activation profile on some tissues. We therefore focused subsequent tauro-BA studies on TDCA, the second-most active AOS ligand identified in this screen. We therefore conducted a concentration-response analysis for 3 bile acid ligands, DCA (0.01–10 µM), TDCA (0.001–1 µM), and GDCA (0.01–10 µM)(Figure 3). We observed a range of sensitivities to these ligands and concentrations. To assist in the identification of VSNs with similar tuning patterns, we performed cluster analysis, which revealed groups of VSNs with similar tuning properties across ligands and concentrations (Fig. 3A).

**Figure 3).**
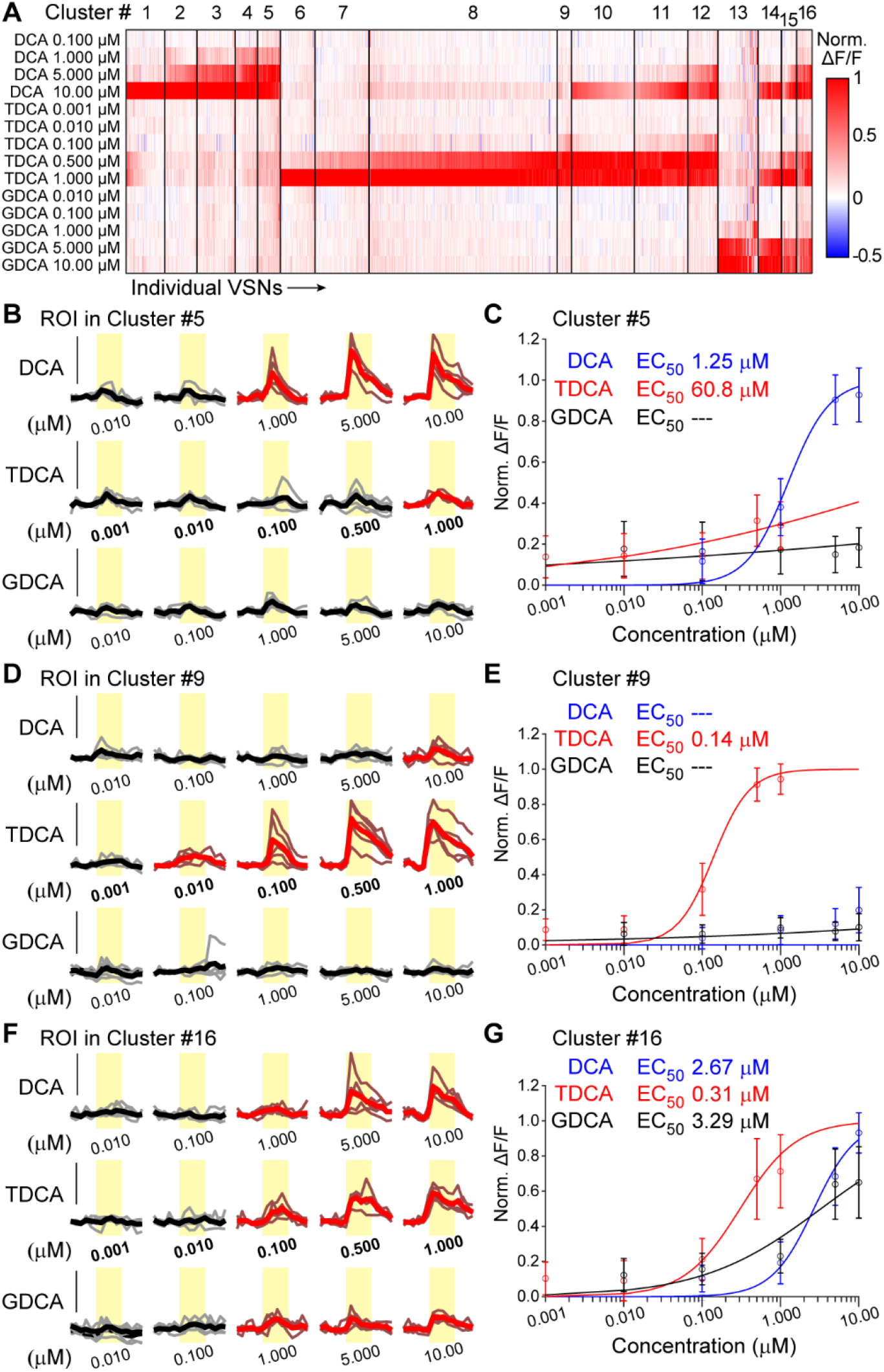
VSNs respond to tauro-BAs at submicromolar concentrations. (**A**) Clustered heat map of 1,857 responsive VSNs exposed to DCA, TDCA, and GDCA at a range of concentrations (7 tissues from 2 males, 2 females). (**B, D, F**) Peri-stimulus time histograms of stimulus responses to ROIs from Cluster 5 (**B**), Cluster 9 (**D**), and Cluster 16 (**F**). Bold traces reflect mean responses to 5 stimulus repeats. Red traces are statistically larger than control (DCA 0.01 µM)(Wilcoxon Rank Sum Test, p<0.01). Yellow rectangles indicate the time of stimulus delivery. Vertical black bars indicate 40% ΔF/F. (**C, E, G**) Concentration-response curves from Cluster 5 (**C**), Cluster 9 (**E**), and Cluster 16 (**G**). Open symbols reflect mean Normalized ΔF/F signals across all cells in the cluster. Error bars reflect standard deviation of the mean. Solid lines reflect median solutions to the Hill Equation (applied to each cell in the cluster, see Methods). EC_50_ values for unresp7onsive ligands are noted as “- - -”.

This analysis identified VSN populations with a high degree of bile acid sensitivity and selectivity. Among these were 418 neurons (22.5% of analyzed cells) that were selective for DCA (Clusters 1-5, Fig. 3). Cluster 5 contained 62 VSNs (3.3% of analyzed cells) with high sensitivity and selectivity for DCA (Fig. 3B-C). VSNs in Cluster 5 had a median EC_50_ value for DCA of 1.25 µM (Fig. 3C). As in the initial screens (Figs. 1-2), many more VSNs responded to TDCA at 1 µM than the other ligands, despite being applied at a 10-fold lower concentration (Fig. 3A). Clusters 6-9 included VSNs that were selectively activated by TDCA (totaling 788 cells, 42.4%). Among these, the 39 VSNs (2.1% of analyzed cells) in Cluster 9 displayed the highest TDCA sensitivity (Fig. 3D-E). VSNs in Cluster 9 had a median EC_50_ value for TDCA of 140 nM (Fig. 3C). We also observed 396 VSNs (21.3% of analyzed cells) with high TDCA sensitivity that were also activated by DCA (Clusters 10-12, Fig. 3A). Among the 81 VSNs (4.4% of analyzed cells) in Cluster 12, the median EC_50_ for TDCA was 126 nM and the median EC_50_ for DCA was 5.64 µM. Cluster 13 contained 109 VSNs (5.9% of analyzed cells) with selective responsiveness to GDCA, with the median EC_50_ for GDCA of 2.17 µM. We had not identified a GDCA-selective population in our initial screen, which utilized 1 µM GDCA. Lastly, we observed 146 VSNs (7.9% of analyzed cells) responded to DCA, TDCA, and GDCA (Clusters 14-16, Fig. 3A). Cluster 16, which contained 42 VSNs (2.3% of analyzed cells), displayed a median EC_50_ value to DCA of 2.67 µM, to TDCA of 310 nM, and to GDCA was 3.29 µM (Fig. 3F-G). These cells resembled those seen in the initial screen (Cluster 9). These results further strengthened our understanding of tauro-BA potency on VSNs, and suggest that TDCA and other tauro-BAs, even if they were present at lower concentrations in the environment, might be potent activators of AOS-mediated behavior.

### Taurine-conjugated bile acids vary with species and microbiome

Innate behaviors pertaining to conspecific and heterospecific interactions are influenced by environmental stimuli sensed by the VNO (Chamero, et al., 2007; Isogai, et al., 2018; Kimchi, et al., 2007; Osakada, et al., 2018; Papes, et al., 2010; Roberts, et al., 2010; Stowers, et al., 2002). Prior molecular analysis of mouse feces indicated that tauro-conjugated bile acids in mouse feces was minimal (Hagey, et al., 2010). However, tauro-BAs were found in the excretions of reptiles (Hofmann, et al., 2010). Other studies have linked the action of gut flora to the complement of bile acids present in feces (Golubeva, et al., 2017). To investigate potential natural sources of tauro-BAs that may be relevant to mice, we performed mass spectrometry analysis on two potential sources of bile acid variability: species and gut microbiome.

We first collected, with the assistance of Herpetology Division of the Dallas Zoo, fecal samples from multiple reptiles, including monitor lizards, snakes, skinks, and tortoises (Table 1). We made aqueous extracts of these samples, and subjected them to mass spectrometry analysis. Consistent with prior studies, tauro-BAs were not detectable in mouse fecal extracts, but were detected in 5 of 8 reptile species. Of note, the tauro-BAs were found evident in species that were fed carnivorous diets, including rats and mice. These data suggest tauro-BAs are components of predator feces with biological activity in the VNO.

We next collected feces from adult male and female germ-free (gnotobiotic) mice and conventionally fed controls, and conducted a simple test of bile acid abundance across unconjugated and conjugated bile acids (Table 2). These results revealed, consistent with prior studies, that CA, DCA, and CDCA are present in conventionally fed mouse feces. There were no detectable levels of tauro-BAs in these conventionally fed mouse fecal extracts (Table 2). In gnotobiotic mouse feces, these common bile acids were undetectable, reflecting a major shift in bile acid content. In addition to the loss of detectable unconjugated bile acids, taurocholic acid (TCA), which was undetectable in conventional, was detected in both male and female mouse fecal extracts at lower concentrations than bile acids detected in conventionally fed mouse feces (Table 2). Combined, the data presented in Tables 1 and 2 confirm that tauro-BAs are increased in other species (specifically, mouse predators), and with alterations of the gut microbiome.

**Table 2.** Mass spectrometry analysis of unconjugated bile acids (CA, DCA, CDCA, LCA), tauro-BAs (TCA, TDCA, TCDCA, TLCA), and glycine-conjugated bile acids (GCA, GDCA, GCDCA, GCLA). All values in µM.

| (in $\mu\text{M}$) | GF$\text{♀}$ | GF$\text{♂}$ | CF$\text{♂}$ | GF$\text{♀}$ |
| --- | --- | --- | --- | --- |
| CA | n.d. | n.d. | 9.75 | 18.70 |
| DCA | n.d. | n.d. | 9.77 | 36.31 |
| CDCA | n.d. | n.d. | 10.13 | 37.18 |
| LCA | n.d. | n.d. | n.d. | n.d. |
| TCA | 2.59 | 1.29 | n.d. | n.d. |
| TDCA | n.d. | n.d. | n.d. | n.d. |
| TCDCA | n.d. | n.d. | n.d. | n.d. |
| TLCA | n.d. | n.d. | n.d. | n.d. |
| GCA | n.d. | n.d. | n.d. | n.d. |
| GDCA | n.d. | n.d. | n.d. | n.d. |
| GCDCA | n.d. | n.d. | n.d. | n.d. |
| GLCA | 0.0151 | 0.0059 | n.d. | n.d. |

### Taurine-conjugated bile acids overlap with gnotobiotic feces responses in the VNO

Given that TCA levels became detectable in gnotobiotic mouse feces, we hypothesized that TCA or some other (possibly unknown) VSN chemosignals may differ between gnotobiotic and conventional feces. We performed OCPI experiments in which we delivered a panel of stimuli comprising 4 unconjugated bile acids 4 tauro-BAs, and conventional and gnotobiotic mouse feces extracts (Fig. 4). As in prior studies, we observed many VSNs that display overlapping sensitivities to conventional mouse feces extracts and common primary and secondary bile acids (Cluster 2)(Fig. 4A, B, E). Interestingly, the responses of VSNs to 1:300 dilutions of conventional mouse feces extracts were almost completely non-overlapping with VSNs that responded to 1:300 dilutions of germ-free mouse feces extracts (Fig. 4E). In contrast, we observed many neurons that responded to 1:300 dilutions of germ-free mouse feces extracts, and were also responsive to tauro-BAs (Cluster 23)(Fig. 4C, D, E). UMAP dimensionality reduction on the tuning curves in this assay revealed broad separation between groups (Fig. 4F). There was a visible divide, with one side of the projection including VSN clusters that responded to conventional mouse feces extracts and unconjugated bile acids, and the other side containing VSN clusters that responded to germ-free mouse feces extracts and tauro-BAs (Fig. 4F). The differences we observed in VSN responsiveness to germ-free and conventional fecal extracts (from both male and female mice) parallels the mass spectrometry analyses (Table 2), and furthermore suggests that more differences in chemosignals remain to be discovered in these conditions.

**Figure 4).**
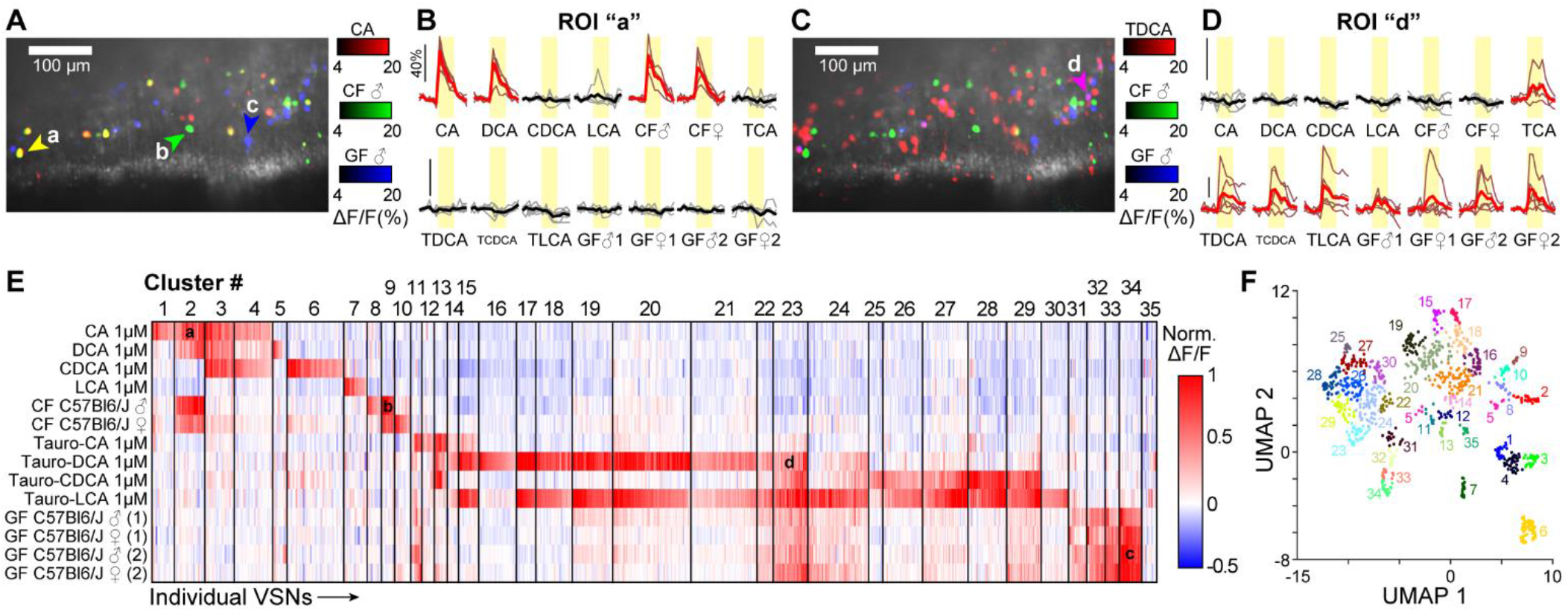
OCPI analysis of unconjugated bile acids, tauro-BAs, conventional mouse feces extracts (CF), and germ-free mouse feces extracts (GF, 1:300 dilutions, samples from both male and female mice). (**A**) Single image frame from a 51-frame stack showing GCaMP6s-expressing VSNs (gray) colorized based on their responsiveness to 1 µM CA (red), conventionally CF (green), GF (blue). (**B**) Peri-stimulus time histograms of the VSN within ROI “a” in Panel A. Red traces indicate p < 0.01 (Wilcoxon rank sum test compared to Ringer’s control, not shown). (**C-D**) Same as Panels A-B but with 1 µM TDCA (red), CF (green), and GF (blue). (**E**) Clustered heat map of ∼1,000 responsive VSNs. Overlaid “a-d” letters refer to VSNs labeled in Panels A-D. (**F**) UMAP (dimensionality reduction) of the tuning properties of these VSNs.

### Taurine-conjugated bile acids induce aversion-related behavioral changes in mice

Given that tauro-BAs are increased in mouse predator feces and germ-free mouse feces – conditions that cause distress or negative health effects – we hypothesized that tauro-BAs may cause aversion in mice. We conducted a series of behavioral assays to assess mouse behavioral responses to tauro-BAs, using TDCA as our representative of the group (Fig. 5). We chose to explore several non-social behavioral contexts in order to ascertain whether (1) TDCA was distinguishable from other BA ligands on its own, and (2) whether TDCA, when added to familiar blends of fecal chemosignals (female mouse feces extract) induces observable changes in behavior. In all tests, we conducted studies on both male and female mice, and observed few sex-specific changes related to tauro-BA detection. We specifically conducted an odor preference test (Zou, et al., 2015), coating porous plastic particles of approximately the same size and shape as mouse feces. During the five-minute test, each mouse was allowed to investigate fake fecal pellets coated with either DCA (50 mg/ml), or TDCA (35 µg or 50 mg), with vehicle-only (water) exposed pellets in the control dish (Fig. 5A-E). DeepLabCut was used to track the position of the animal, and the petri dishes themselves (Mathis, et al., 2018). In these conditions, with no other odorants present in the environment, mice displayed mild preference towards DCA-coated particles, and mild aversion to TDCA-coated particles (Fig. 5B-C, 2-way ANOVA F(2,101) = 9.31, p < 0.0004, main effect of stimulus). In experiments in which DCA was present, both male and female animals traveled less overall distance compared to TDCA (Fig 5D-E, 2-way ANOVA (F2,50) = 5.34, p < 0.008, main effect of stimulus). Thus, in a context in which mice experience feces-like particles and monomolecular tauro-BAs, they display mild aversion.

**Figure 5).**
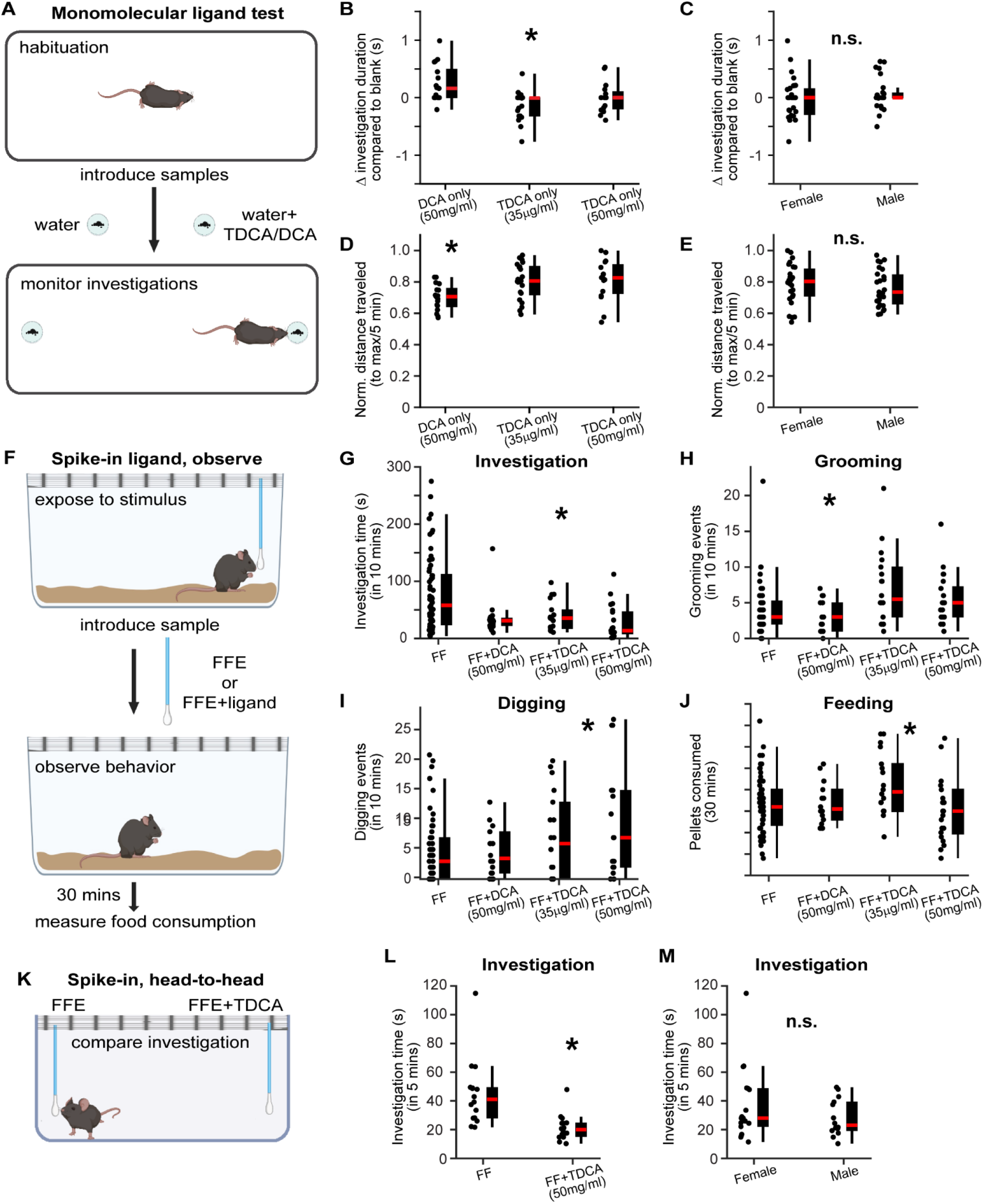
Mice respond to tauro-BAs with aversion-associated behaviors. (**A**) Setup of two-choice assay of fecal particles coated with DCA (50 mg/ml) or TDCA (35 µg/ml and 50 mg/ml). Animals were tracked using DeepLabCut and their interactions with either sample dish scored. (**B-C**) Investigation time relative to negative control (water). (**D-E**) Total distance traveled during sample tests. (F) Setup of spike-in test for DCA and TDCA (same concentrations used in (**A**). (**G-J**) Results of behavior tests and feeding behavior 30 mins following the exposure. (**K**) Setup of two-choice spike-in experiment. (**L-M**) Dif1fe1rence in investigation time. Asterisks indicate p < 0.05 (ANOVA [A-E], Linear mixed effects model [F-M]). “n.s.” : Not statistically significant.

Given that mice do not naturally encounter isolated individual bile acids, we next sought to determine how TDCA affected behaviors towards familiar conspecific blends of bile acids (female mouse feces extracts, Fig. 5F-J). We exposed mice to cotton swabs dipped in mouse feces extract or mouse feces extract supplemented with DCA or TDCA (same concentrations as above) and assessed several behaviors, including direct investigation time (Fig. 5G), self-grooming (Fig. 5H), and digging (Fig. 5I). Because previous studies indicated that tauro-BAs were associated with anorexia-like behavior (Perino, et al., 2021), following these exposures we assessed mouse feeding behavior by giving them 30 minutes of *ad libitum* access to small chow pellets, and measuring the number consumed (Fig. 5J). We observed decreased investigation time in these “spike-in” experiments compared to female feces extracts alone (Fig. 5G, linear mixed effects model; F(3,106) = 7.4, p < 0.0001, main effect of stimulus). Predatory pheromones such as snake odor regulate the grooming and digging behavior of mice (Dell’Omo, et al., 1994). Mice expressed more self-grooming (Fig. 5H, linear mixed effects model F(3,106) = 4.7, p < 0.004, main effect of stimulus), and digging behaviors when TDCA was spiked-in compared to female feces alone or when female feces was spiked with DCA (Fig. 5I, linear mixed effect model, F(3,106) = 2.34, p < 0.077, pairwise post-hoc comparison between TDCA 50 mg/ml and other conditions, p < 0.05). Feeding behavior was mildly decreased in the TDCA (35 µg/ml) condition (Fig. 5J, linear mixed effect model; F(3,106) = 2.93, p < 0.037, main effect of treatment). For the feeding behavior, we observed a main effect of sex, with females consuming more pellets than males (linear mixed effects model, F(1,106) = 10.1, p < 0.0019). Finally, we performed a second two-choice assay, this time using two cotton swabs spiked with female feces extracts alone and female feces extracts spiked with TDCA (50 mg/ml, Fig. 5K-M). In this simple context, investigation time of the swab with spiked-in TDCA was markedly decreased (linear mixed effect model; F(1,29) = 15.0, p < 0.0006, main effect of stimulus). Collectively, these assays provide evidence that mice behave differently when tauro-BAs are sensed in the environment, and that tauro-BAs elicit mildly aversive behaviors.

## Discussion

The lack of knowledge about the full complement of natural AOS ligands (and corresponding receptors) is a major barrier towards understanding the behavioral and physiological impacts of the accessory olfactory system. In this study, we identified taurine-conjugated bile acids as a new class of AOS ligands with remarkable potency. The participation of bile acids in social chemosensation has been noted for many years in fish, where these molecules serve as pheromones (Buchinger, et al., 2014; Cong, Zheng, Ren, Chéron, et al., 2019; Cong, Zheng, Ren, Cheron, et al., 2019). Our previous work showed that fecal bile acids function as natural AOS ligands in mice, indicating that bile acid chemosensation is conserved between fish and mammals (Doyle, et al., 2016). In the context of our growing knowledge of bile acids as AOS ligands, the addition of this new group of potent AOS ligands will help to advance our understanding of the AOS and its many impacts on mammalian social and reproductive behaviors.

### Taurine-conjugated bile acids are potent AOS ligands

Using the OCPI platform, which allows the simultaneous monitoring of hundreds to thousands of VSNs per tissue (Holekamp, et al., 2008; Turaga, et al., 2012), we screened through commercially available bile acid ligands and discovered that tauro-BAs activate a larger VSN population than unconjugated and other conjugated bile acids, and do so at lower concentrations that prior screens (Fig. 1)(Doyle, et al., 2016; Nodari, et al., 2008). Most tauro-BA-sensitive VSNs were unresponsive to other bile acids and sulfated steroid ligands, even at higher concentrations (Figs 2-3). Because of the strong links between chemosensory tuning and chemosensory expression in peripheral sensory neurons (Lee, et al., 2019; Wong, et al., 2020), these results strongly suggest that tauro-BAs are sensed by a set of receptors without an identified natural ligand. Previous studies indicated that multiple V1R-family receptors, including members of the V1rc and V1re clades, are bile acid-sensitive. It is possible that other receptors in these clades also sense tauro-BAs. However, given the lack of direct links between receptor clade and ligand sensitivity in the VNO (Lee, et al., 2019), it may be that receptors from other clades participate in tauro-BA detection.

VSN sensitivity and selectivity for tauro-BAs is lower than most of the other major known AOS ligand classes. There are many hundreds of naturally occurring bile acids, conjugates, and variants (Hofmann, et al., 2010), and there are many left to evaluate in the future. The detection of these signals by peripheral vomeronasal neurons, their integration by downstream neurons in the accessory olfactory bulb, and their influence on neural circuits controlling animal behavior remain unknown. The combinatorial nature of chemosensory was on clear display in this study, and each molecule added to the list of known AOS ligands advances our understanding of this important chemosensory pathway.

### Taurine-conjugated bile acids vary with species and gut microbiome

The natural conditions under in which mice encounter bile acids is apparent when considering bile acids shed by mice. Fecal particles are durable in the environment, even in laboratory settings, where cage changes associated with mouse husbandry are spaced days-to-weeks apart. Fecal particles dry rapidly in the environment, concentrating molecules within, and bile acids are very stable in these conditions (Buchinger, et al., 2014; Feller, et al., 2021). The striking activity caused by tauro-BAs in the VNO suggested that mice have evolved receptors for these ligands to support some important behavioral function. Because taurine-conjugated bile acids have been reported in reptiles, we sought to determine directly whether any of the tauro-BAs active on VSNs were present in reptile feces extracts (Table 1). Confirmation that TCA, TDCA, TCDCA, and TLCA were detectable in carnivore lizards, many of which were fed rodent diets in captivity, suggests at least one purpose of VNO sensitivity to tauro-BAs: predator detection and avoidance. It is worth noting that similar reptile fecal samples to those included in this analysis caused patterns of threat assessment behavior in mice (Wang, et al., 2023), suggesting that some of the molecules that contributed to these behavioral observations might be tauro-BAs.

The importance of the gut microbiome on an organism’s complement of bile acids is well-recognized (Lin, et al., 2024). Because mice evolved in conditions where food sources were highly variable, and variably scarce, gut flora are likely to change dramatically across the life of each animal. Rather than explore the dizzying number of variables associated with diet and gut microbiome effects on bile acids, we simply sought to determine whether any of the tauro-BAs that activate the VNO vary with complete removal of the gut microbiome (Table 2). Consistent with prior work, we found that conventional mouse feces contained multiple primary and secondary unconjugated bile acids, including CA, DCA, and CDCA (Doyle, et al., 2016). It was somewhat surprising that CA and CDCA, primary bile acids that are generated from cholesterol in the liver (i.e. not requiring gut microbes for their generation), were undetectable in germ-free male and female mouse feces (Table 2). None of the analyzed tauro-BAs were detectable in conventional male or female mouse feces, but TCA was reliably detected in germ-free mouse feces, confirming that tauro-BAs vary with this important biological variable.

The evidence that gut microbiome alters tauro-BA levels suggested the capacity for VSNs to discriminate between germ free and conventional mouse feces, which we investigated and confirmed using the OCPI microscopy platform (Fig. 4). This experiment confirmed the association between VSN sensitivity to conspecific fecal extracts and unconjugated bile acids (Doyle, et al., 2016). Comparing VSN responsiveness to germ-free and conventional mouse feces extracts revealed stark separation between populations sensitive to each (Fig. 4). We observed almost no overlap between populations of VSNs that responded to conventional fecal extracts and those that responded to germ-free fecal extracts. Among the populations of VSNs that responded to germ-free mouse fecal extracts, we observed several that responded to tauro-BAs at 1 µM (Fig. 4C, D, E). These results confirm the capacity for VSNs to (1) discriminate between germ-free and conventional mouse feces and (2) to associate tauro-BAs with alterations in the gut microbiome. It is worth noting that there were many neurons that were sensitive to germ-free mouse feces but unresponsive to any other ligand in our stimulus panel in this experiment, suggesting that there are many, potentially unknown, new AOS ligands to discover in gnotobiotic mouse feces. Overall, this relatively straightforward experiment provided valuable insight into links between the gut microbiome and tauro-BAs, and represents an initial foray into the rich, interwoven nature of diet, microbiome, and other biological variables that may affect bile acid chemosensation.

### Mice respond to TDCA in the environment with aversive behaviors

The potency of tauro-BAs suggests it activates the AOS at lower concentrations than other ligands, including unconjugated bile acids. The association between tauro-BAs and predator feces and loss of gut flora suggests that tauro-BA detection might be aversive to mice, even in a non-social, non-threatening laboratory context. We performed laboratory behavioral tests using TDCA, the second-most active tauro-BA, to avoid potential toxicity effects of TLCA, the most-active tauro-BA identified in our screen. We designed these experiments to investigate mouse behavior to monomolecular TDCA compared to DCA. In this context, TDCA serves as a novel ligand, as it would unlikely have been encountered by the mouse in conventional housing conditions. Mice showed mild avoidance to TDCA itself when presented on an inert plastic particle resembling feces, but mild preference for the familiar DCA (Fig. 5A-E). We then took a “spike-in” approach to test the effect of the unfamiliar TDCA when it was added to a familiar blend of bile acids, with extra DCA, which is already present in conventionally raised female mouse feces (Table 2)(Doyle, et al., 2016) serving as a control (Fig. 5F-J). In these tests, we observed reduced investigation time, increased grooming events, increased digging events, and increases in feeding behavior. In all these cases, the effects were mild, though significant. This is perhaps unsurprising, as mice are unlikely to have experienced any of other natural aspects of TDCA sensation, which would involve other chemosignals and multisensory inputs (e.g. if encountered in the context of a predator or sick mouse). Nevertheless, a direct comparison between familiar female mouse feces extract and the fecal extract with spiked-in TDCA showed a clear aversion, despite the lack of a prior learned association. Together, these results indicate that detection of tauro-BAs leads to behavioral aversion, supporting a potential role for tauro-BAs in steering mice towards conspecifics with healthy gut microbiomes.

In summary, we have demonstrated that taurine-conjugated bile acids are AOS ligands with the highest potency yet observed in this molecular class. Tauro-BAs vary with species and gut microbiome, increasing in the feces of mouse predators and mice lacking a gut microbiome. A representative of the tauro-BAs, tauro-deoxycholic acid, elicited mildly aversive responses in mice with no prior training or association. These results establish new and deeper links between gut physiology, inter-species communication and bile acid chemosensation.

## MATERIALS AND METHODS

### Animals

All animal experiments were performed in accordance with the Institutional Animal Care and Use Committee at the University of Rochester Medical Center and follow the guidelines of the National Institutes of Health. Physiological experiments were performed with OMP^tm4(cre)Mom/J^ knock-in mice (OMP-Cre mice; Jackson Laboratory stock number 006668) mated with Gt(ROSA)^26Sortm96(CAG-GCaMP6s)Hze/J^ mice (Ai96 mice); the Jackson Laboratory order no. 024106). These mice express the genetically encoded Ca^2+^ indicator GCaMP6s (OMP-Cre^+/−^, Ai96^+/−^) in VSNs, hereinafter referred to as OMPxAi96 mice. All experiments were performed on adult mice (age 8–16 weeks). 2–5 animals per cage were maintained on a 12-hour light/dark cycle and had *ad libitum* access to food and water. The number, strain, and sex of animals used in each experiment are described in the corresponding figure legends.

### Solutions and stimulus presentation

Bile acid (BA) stimuli included CA, DCA, CDCA, LCA, DHCA, UDCA, β-MCA, λ-MCA, δ-MCA, TCA, TDCA, TLCA, TCDCA, GCA, GDCA, GLCA, GCDCA, Iso-DCA, 7-KetoDCA, 7-KetoLCA and 12-KetoDCA. Sulfated steroids (SS) included epitestosterone-17-sulfate (A6940), 17α-estradiol-3-sulfate (E0893), 5α-pregnane-3β-ol-20-one sulfate (P3865), and corticosterone-21-sulfate (Q1570). BA and SS are purchased from Sigma (St. Louis, MO), Steraloids Inc. (Newport, RI, USA) and MedChemExpress (Monmouth Junction, New Jersey). Stock solutions (20 mM) of all BAs and sulfated steroids were prepared in methanol and diluted to their final concentration in Ringer’s solution containing 115 mM NaCl, 5 mM KCl, 2 mM CaCl_2_, 2 mM MgCl_2_, 25 mM NaHCO_3_, 10 mM HEPES. and 10 mM glucose. All sulfated steroids were diluted to 10 μM (1:2000) for experiments. BAs were diluted to a concentration range of 0.001 to 10 μM. Control stimuli consisted of Ringer’s solution containing 1:2000 methanol (the highest methanol concentration in any individual stimulus). Stimuli were applied for 15 seconds using an air pressure-operated reservoir via a 16-in-1 multi-barrel “perfusion pen” (AutoMate Scientific, Berkeley, CA, USA).

For behavioral experiments, 50 mg/ml DCA or TDCA was prepared in 1 ml female feces extract (FF). Exposure to TDCA, DCA, or FF was achieved by dipping them in fake fecal pellets or by using cotton swabs. Fecal extracts from female mice were prepared by dissolving 5 g of feces in 50 ml of distilled water (dH_2_O). After the fecal mixture was stirred for two minutes, it was placed on ice and shaken continuously throughout the night. Following a 2-minute vortex on the second day, the feces mixture was centrifuged twice (10 min at 2400 × g at 4°C and 30 min at 2800 × g at 4°C) to homogenize it. Using a 0.22 μm filter (“s_filtered”), the supernatant from two centrifugations was combined and placed in collection tubes. The “solid” form of feces was also preserved at −80°C. The control stimulus, or “control,” was pure water (dH_2_O) (Wang, et al., 2023).

### Volumetric VNO Ca^2+^ imaging

Similar to previous studies (Turaga, et al., 2012; Wong, et al., 2020; Wong, et al., 2018), VNOs were dissected and the vomeronasal epithelium was carefully removed under a dissection microscope (Leica Microsystems, Buffalo Grove, IL, USA) after deep isofluorane anesthesia and rapid decapitation. The vomeronasal epithelium was placed in a specially designed imaging chamber after being mounted on nitrocellulose paper (Thermo Fisher Scientific, Atlanta, GA, USA). A custom-made OCPI microscope was used to perform volumetric Ca^2+^ imaging with previously reported refinements (Wong, et al., 2020; Wong, et al., 2018). Utilizing custom software that coordinated imaging and stimulus delivery through a randomized, interleaved stimulus delivery system (AutoMate Scientific), the fluorescence of GCaMP6s was measured. Approximately 700 μm laterally, 250 to 400 μm axially, and ∼150 μm in depth were covered by image stacks with 51 frames, acquired once every 3 s (∼0.33 Hz). Five consecutive stacks (∼15 s) of each stimulus were given, with at least 10 stacks (≥30 s) elapsed between stimulus trials. Three full randomized, interleaved stimulus blocks were completed in each of the experiments that were examined.

### Data analysis of volumetric VNO Ca^2+^ imaging

Custom MATLAB software was used for data analysis similar to previous studies (Hammen, et al., 2014; Turaga, et al., 2012). Rigid registration was applied to image stacks, and then they underwent nonrigid warping. Secondly, the mean voxel intensity in three successive pre-stimulus stacks was subtracted from the mean voxel intensity of three stacks during stimulus delivery, and the result was divided by the mean pre-stimulus intensity value to determine ΔF/F, the relative change in GCaMP6s intensity. The process involved manually drawing volumetric ROIs around the cell bodies of spontaneously active neurons and well-registered VSNs that exhibited consistent response to stimulation. A matrix of fluorescence intensity was produced by calculating the mean voxel intensity for each ROI after the volumetric ROI selection was applied to each of the approximate 1200 image stacks in the experiment. This matrix is used to calculate the across-trial mean ΔF/F for all ROIs and stimulus applications. In order to account for potential effects of valve switching, the across-trial Ringer’s control stimulus was compared to the ΔF/F of each ROI that was evoked by the stimulus. This method measured the stimulus responsiveness. The ROIs exhibiting a positive ΔF/F response and a P < 0.05 to a stimulus were deemed responsive, and the nonparametric Wilcoxon-Mann-Whitney test was utilized to determine a P-value linked to the stimulus response. Clustering was performed using density-based merging (DMB), part of the “runUMAP” package available in MATLAB. DBM parameters were selected based on manual inspection of results, and inspection of pairwise linear discriminant analysis (LDA), which was used to evaluate cluster separation via the discriminability index statistic (d′, see Fig. 2F). Satisfactory cluster separation was confirmed by a discriminability index score greater than 1.64 (when the z-score equivalent P value is equal to 0.05).

### Mass spectrometry analysis

Reptile fecal samples were collected by members of the Division of Herpetology at the Dallas Zoo as part of routine cage maintenance and cleaning, and did not involve any alteration to standard animal housing routines. Fresh fecal samples were placed into plastic tubes and stored at −20 °C until collection by laboratory staff. Aqueous extracts from these samples were collected as noted above (Solutions and stimulus presentation), and aliquots were stored at −80 °C until use. Mass spectrometry analysis of bile acids was performed similar to previous methods (Han, et al., 2015). Briefly, verified monomolecular bile acid standards purchased from Steraloids, Santa Cruz Biotechnologies (Santa Cruz, CA, USA), Toronto Research Chemicals (Toronto, Ontario, Canada), or C/D/N Isotopes (Pointe-Claire, Quebec, Canada) and used as isotope-labeled standards. Samples were analyzed via ultrahigh performance liquid chromatography/multiple-reaction monitoring-mass spectrometry (UPLC-MRM-MS). This involved an Ultimate 3000 RSLC system (Dionex Inc., Amsterdam, The Netherlands) coupled to a 4000 QTRAP mass spectrometer (AB Sciex, Concord, ON, Canada) via a Turbo Ionspray electrospray ionization (ESI) source, which was operated in the negative ion mode. A BEH C18 (2.1 mm × 150 mm, 1.7 μm) UPLC column (Waters Inc., Milford, MA) was used for the gradient elution, with 0.01% formic acid in water (solvent A) and 0.01% formic acid in acetonitrile (solvent B) as the mobile phase. The collision energy for each group-specific MRM transition was the median of the collision energies for the same transition for all the isomeric BAs in each group.

Germ-free mouse fecal extracts were obtained from adult male and female C57Bl6 mice housed in a gnotobiotic housing facility via University of Rochester’s Division of Comparative Medicine as part of routine cage maintenance. Aqueous extracts of germ-free fecal samples, as well as from C57Bl6 mice maintained in a standard/conventional mouse housing facility, were diluted in dH_2_O and submitted for analysis via Creative Proteomics (Shirley, NY, USA). Briefly, samples were analyzed via an AB SCIEX Qtrap 5500 mass spectrometer operated in negative ion mode, and connected to a Waters ACQUITY Ultra Performance Liquid Chromatographer. Samples were run over a Waters ACQUIRT UPLC BEH C18 column (2.1 mm × 100 mm, 1.7 μm) for the gradient elution, with 0.05% formic acid in water (solvent A) and 0.05% formic acid in acetonitrile (solvent B) as the mobile phase.

### Behavioral test setup

All behavioral tests were performed under dim red light during the dark cycle. Mice were solo housed for 10 days before the start of experiment. In the TDCA preference test, the experimental animals are placed in fresh cages that are provided with food and water so that they can acclimatize in the behavioral room for at least two hours. Mice are habituated in large, open experimental cages for fifteen minutes. One hour before the experiment, ellipse-shaped, one-centimeter-long 3D-printed fake fecal particles were submerged in TDCA, FF, or water. n = 5–7 fake fecal particles were placed in the Petri dish and kept moist for the duration of the experiment. One Petri dish was used exclusively for one odor in order to prevent cross-contamination. Petri dishes containing fecal pellets were positioned in the two corners of the cage and a five-minute video recording was made. After completion of the investigation experiment, mice were allowed to be exposed to FF or TDCA pellets for an additional half hour. The nesting material was added after the Petri plates were removed, and the nesting behavior was watched for three hours. To avoid positional bias, the position of the Petri plates was swapped for each presentation.

For the TDCA investigation assay, mice were deprived of food for 12 hours before the start of the experiment. Five minutes after being exposed to FF or TDCA, mice were able to access food pellets. Habituation time in the experimental cage was 65 minutes. A cotton swab dipped in FF or TDCA was placed through the cage bar and the investigation was captured on video for ten minutes. FFMPEG was used to convert videos that were recorded at 30 frames per second (FPS) into frames. An additional double-blind analysis was performed for these videos. The number of frames that showed the mice examining the cotton swab was counted manually, and the number of investigation events and investigation times were scored. A similar process was used to manually annotate and score the number of frames corresponding to digging and grooming. The cotton swab was removed after ten minutes, and after five minutes forty food pellets were placed to a Petri plate to measure feeding behavior after thirty minutes. We finally elucidated the chemosensory functions of TDCA, by designing a mouse behavioral experiment in which mice is allowed to investigate cotton buds having either FF extract or FF spiked in with TDCA. The experiment was conducted for 10 min and the time investigating the cotton buds were manually calculated as investigation time. TDCA investigation assays using cotton swabs were scored manually by a blinded individual.

### DeepLabCut tracking

TDCA preference test videos in the fake fecal pellets assay (Fig. 5A-E) were tracked using DeepLabCut (2.2.0.3) (Mathis, et al., 2018; Nath, et al., 2019). The DeepLabCut model included seven labels in the mouse body (nose, left ear, right ear, base of tail, mid_1 (between the left and right ear), mid_2 (center of the posterior body), mid_3 (between mid_1 and base of the tail). To track the petri dishes, which occasionally moved as a result of mouse inspection, we trained the model to track eight points around the circumference of each petri dish. There were minor variations in the experimental conditions such as lighting, testing arena, animal and background, which required the iterative addition of new videos and labeled frames to create a robust and accurate model. Models were trained for 200,000-800,000 iterations using the “ResNet-50” neural network on more than 1000 labeled frames with default parameters in the DeepLabCut GUI. On assessment, eighty percent of labeled frames were used to training the network, and the other twenty percent were used for network assessment. We proceeded to analyze videos in which if the test error with p-cutoff was approximately five pixels (image size: 1160 × 716 pixels; 1 mm ≈ 1 pixels). To smooth over random/spurious poorly tracked frames, we applied a rolling median filter with n = 7 frames to the raw tracking data. Accuracy was manually verified in DeepLabCut-generated labeled videos before they were finalized.

## Acknowledgments

We thank members of the Chemosensation and Social Learning Laboratory for helpful feedback. We especially thank Michael Mastrangelo for critical animal husbandry support. We thank the Dallas Zoo Department of Herpetology for providing reptile fecal samples (DZ Project S2019-2), and the laboratory of Dr. Felix Yarovinsky for providing gnotobiotic fecal samples.

## Author contributions

V.H. and J.P.M. designed project, analyzed data, and wrote the manuscript. V.H. performed microscopy experiments. V.H., M.M., and J.P.M. analyzed microscopy data. V.H. and L.R. performed behavior experiments. V.H. and J.P.M. analyzed behavioral data. J.W. and J.G.M collected reptile samples and conducted mass spectrometry experiments.

